# Tire abrasion particles negatively affect plant growth even at low concentrations and alter soil biogeochemical cycling

**DOI:** 10.1101/2021.04.19.440468

**Authors:** Eva F Leifheit, Hanna L Kissener, Erik Faltin, Masahiro Ryo, Matthias C Rillig

## Abstract

Tire particles (TPs) are a major source of microplastic on land, and considering their chemical composition, they represent a potential hazard for the terrestrial environment. We studied the effects of TPs at environmentally relevant concentrations along a wide concentration gradient (0 – 160 mg g^−1^) and tested the effects on plant growth, soil pH and the key ecosystem process of litter decomposition and soil respiration. The addition of TPs negatively affected shoot and root growth already at low concentrations. Tea litter decomposition slightly increased with lower additions of TPs but decreased later on. Soil pH increased until a TP concentration of 80 mg kg-1 and leveled off afterwards. Soil respiration clearly increased with increasing concentration of added TPs. Plant growth was likely reduced with starting contamination and stopped when contamination reached a certain level in the soil. The presence of TPs altered a number of biogeochemical soil parameters that can have further effects on plant performance. Considering the quantities of yearly produced TPs, their persistence, and toxic potential, we assume that these particles will eventually have a significant impact on terrestrial ecosystems.

## 1. Introduction

Microplastic pollution has become a global threat to oceans, rivers and soils (Blettler et al., 2018; Haward, 2018; Rillig and Lehmann, 2020). The emerging research on terrestrial ecosystems has been focused on microplastic originating from littering, sewage sludge or mulching. However, a major source of microplastic particles on land are tire particles (TP) originating from abrasion of tires on roads (Baensch-Baltruschat et al., 2020; Evangeliou et al., 2020). TPs are considered microplastic as they contain a large part of synthetic polymers, have a solid state, are insoluble in water, and have particle size ranges typical of microplastic (1-1000 μm) (Hartmann et al., 2019). Kole et al. (2017) estimate the global annual emission of particles from tires to be more than 3.3 megatons. Although partially retained by separate sewer systems in urban areas, approximately two thirds of these particles end up in soils via runoff or transport by air (Panko et al., 2013). Amounts of TPs in the soil vary a lot: depending on the distance from the road, transport dynamics and the traffic frequency, concentrations can range from 0.1 – 117 g kg^−1^ (Baensch-Baltruschat et al., 2020; Wagner et al., 2018; Wik and Dave, 2009).

Considering the composition of the tires, they represent a potential hazard for the terrestrial environment. Tires usually contain 40-50 % rubber (natural and synthetic), 30-35 % fillers (typically carbon black, silica and chalk), 15 % softeners (e.g. oils), 2-5 % vulcanization agents (S and ZnO) and further additives (Baensch-Baltruschat et al., 2020; Brandsma et al., 2019). Additionally, tires can release other contaminants such as heavy metals and polycyclic aromatic hydrocarbons (PAHs) into the environment (Kreider et al., 2010). Considering these numbers, it is surprising that the effects of TPs in the terrestrial environment have not received more attention so far.

Microplastics often have positive effects on plant performance (de Souza Machado et al., 2018; Lehmann et al., 2020), but also negative effects can occur (Qi et al., 2018; van Kleunen et al., 2020). Tire particles have been tested in past studies for their suitability as potting substrate for plants. The amendment with TPs unambiguously induced morphological and anatomic alterations, as well as growth limitations in the aboveground biomass and altered nutrient concentrations in plant tissue (Bowman, 1994; Schulz, 1987). The authors identified the toxicity of zinc (Zn) as the main problem. More recent studies shifted towards a focus on ecotoxicology, mainly in aquatic ecosystems. Here, toxic effects of TP leachates were connected with Zn, other heavy metals and organic components such as phthalates, resin acids or benzothiazoles (Baensch-Baltruschat et al., 2020).

The impact of TPs on soil biota and soil functions relevant to plant performance is largely unknown. Microplastic in general can affect the growth, metabolism, reproduction and mortality of a wide range of the soil fauna, especially at high concentrations and small particle sizes (Büks et al., 2020). A number of soil functions, such as decomposition or soil aggregation, as well as microbial activity can be altered by microplastics (de Souza Machado et al., 2019). One recent study in a soil ecosystem looked at the effect of TPs on earthworms, showing altered gut microbiota of the animals and impaired survival and reproduction rates (Ding et al., 2020). Not only the particles themselves can be responsible for the observed effects but also the additives contained in the materials (Baensch-Baltruschat et al., 2020). To our knowledge there is no empirical study showing the effect of TPs on plant growth and soil biogeochemical cycling at environmentally relevant concentrations along a wide concentration gradient. Studies used either very high concentrations, such as for potting substrates with up to 66% of the substrate (Bowman, 1994), or used rather low concentrations of ≤ 30 mg g^−1^ (Ding et al., 2020; Schulz, 1987). In this study, we cover a response surface between 0 and 160 mg g^−1^, with the latter being a realistic maximum concentration for TPs in German soils (Baensch-Baltruschat et al., 2020). We performed this study in the light of increasing global traffic volume and thus increased tire wear emissions and the lack of knowledge on the effect of TPs on the terrestrial environment. Given the significant concentrations of toxic components in the TPs we expect negative effects on plant performance, while at the same time carbon compounds and a reduced bulk density will lead to stimulating effects on microorganisms.

## 2. Materials and Methods

We performed a pot experiment with natural soil and a plant, using a concentration gradient of TP pollution from 0 – 160 mg g^−1^ soil with 3 replicates per concentration level. We tested the effects of TPs on plant growth, soil pH and the key ecosystem process of litter decomposition and soil respiration.

### 2.1. Soil and tire particles

For the experiment, a soil-sand mix was prepared. The soil was collected at a field site in Berlin and was classified as Albic Luvisol following FAO classification (loamy sand, 6.9 mg P 100g^−1^ soil (calcium-acetate-lactate), 0.12% N (total), 1.87% C (total), see Leifheit et al. (2015), 56 μm Zn g^−1^ soil (aqua regia digestion)). The air-dried soil was sieved with a 2 mm sieve and mixed with autoclaved sand (50:50 w/w). We diluted the soil with sand in order to facilitate the harvest of the belowground material. The pH of the soil sand mixture was 5.41 (EN 15933:2012). The experimental system was established in white polypropylene pots (Ray Leach Conetainers, Stuewe and Sons Inc.) with a cotton wadding at the bottom and initially filled with 70 g autoclaved sand.

We produced tire abrasion particles from a used car tire (Goodyear M+S 195/60R15 88H) using a portable belt grinding machine (Bosch PBS 75 AE). The particles had an average size of 125 μm within a range of 34-265 μm (determined by photographic analysis, see Fig. S1). Heavy metals in the tire particles were determined with an aqua regia digestion: Cr 175±7 μg g^−1^, Pb 357±29 μg g^−1^, Zn 5089±40 μg g^−1^, Ni 95±3 μg g^−1^ and Cu 453±10 μg g^−1^ (ICP-OES Perkin Elmer Optima 2100DV). TPs typically have a density of 1.13-1.16 g cm^−3^ (Rhodes et al., 2012).

The concentration of microplastic was increased in steps of 10 mg g^−1^ soil, starting at 0 mg TPs g^−1^ soil-sand-mix, up to 160 mg g^−1^, giving 17 different concentration levels with 3 replicate pots per level, summing up to 51 pots in total (see Table 1 for details on added amounts of TPs and the corresponding added heavy metal concentrations). The TP concentrations were chosen based on literature values, using current environmentally relevant concentrations and prospective concentrations of accumulating TP in soil for Germany (Baensch-Baltruschat et al., 2020; Klöckner et al., 2019).

**Table 1:** Amount of tire particles (TP) and the corresponding concentrations of heavy metals that were added to the experimental soil.

| TP<br>(mg g<sup>-1</sup>) | Corresponding concentrations of heavy metals |  |  |  |  | Final Zn<sup>a</sup><br>(mg kg<sup>-1</sup>) |
| --- | --- | --- | --- | --- | --- | --- |
|  | Zn (mg kg<sup>-1</sup>) | Pb (mg kg<sup>-1</sup>) | Cr (mg kg<sup>-1</sup>) | Ni (mg kg<sup>-1</sup>) | Cu (mg kg<sup>-1</sup>) |  |
| 0 | 0 | 0 | 0 | 0 | 0 | 56 |
| 10 | 50.9 | 3.6 | 1.8 | 1 | 4.5 | 106.9 |
| 20 | 101.8 | 7.1 | 3.5 | 1.9 | 9.1 | 157.8 |
| 30 | 152.7 | 10.7 | 5.3 | 2.9 | 13.6 | 208.7 |
| 40 | 203.6 | 14.3 | 7 | 3.8 | 18.1 | 259.6 |
| 50 | 254.5 | 17.9 | 8.8 | 4.8 | 22.7 | 310.5 |
| 60 | 305.3 | 21.4 | 10.5 | 5.7 | 27.2 | 361.3 |
| 70 | 356.2 | 25.0 | 12.3 | 6.7 | 31.7 | 412.2 |
| 80 | 407.1 | 28.6 | 14 | 7.6 | 36.2 | 463.1 |
| 90 | 458.0 | 32.1 | 15.8 | 8.6 | 40.8 | 514.0 |
| 100 | 508.9 | 35.7 | 17.5 | 9.5 | 45.3 | 564.9 |
| 110 | 559.8 | 39.3 | 19.3 | 10.5 | 49.8 | 615.8 |
| 120 | 610.7 | 42.8 | 21 | 11.4 | 54.4 | 666.7 |
| 130 | 661.6 | 46.4 | 22.8 | 12.4 | 58.9 | 717.6 |
| 140 | 712.5 | 50.0 | 24.5 | 13.3 | 63.4 | 768.5 |
| 150 | 763.4 | 53.6 | 26.3 | 14.3 | 68.0 | 819.4 |
| 160 | 814.2 | 57.1 | 28 | 15.2 | 72.5 | 870.2 |

<sup>a</sup>The background concentration of Zn in the experimental soil was 56 µg g<sup>-1</sup>. ‘Final Zn’ represents the final concentration of Zn in the experimental soil, added from TPs plus the background concentration in the soil.

### 2.2. Litter and plant

For the analysis of microplastic effects on litter decomposition we produced mesh bags with a size of 3×2.5 cm containing 300±10 mg of tea leaf litter (Lipton Green Tea, Sencha Exclusive Selection) (see Lehmann et al., 2020).

Seeds of *Allium porrum* (variety “De Carentan”, Floraself) were sterilized with a 0.5% solution of NaClO-bleach for 10 minutes and 70% ethanol for 40 seconds. Sterilized seeds were washed with demineralized water and germinated in autoclaved sand. Seedlings were watered with demineralized water.

### 2.3. Experimental set-up

The experiment was set up in a randomized manner: a labelled pot was picked at random and filled with the soil-microplastic mixture. For each pot, the tire particles were mixed manually with 50 gram of soil-sand-mix in a glass beaker by stirring with a metal spatula. The mesh bag was placed in the centre of the pot when half of the soil-microplastic-mix was added and finally the rest of the soil-microplastic-mix was filled into the pot.

The growth substrate (the soil-microplastic mix) was watered to 60% water holding capacity (WHC) and this soil moisture was kept constant during the experiment by gravimetric watering with deionised water for all pots.

Seedlings of similar height were selected and placed in the centre of the pot. Pots were placed in a randomised manner on two trays and randomised again once per week. Greenhouse conditions were 22/18 °C (day/night) with a daylight period of 12 hours, illuminance: 50 klx, and a relative humidity ~40%. Plants were grown for 6 weeks and the experiment was harvested destructively.

### 2.4. Harvest

The harvest was done in a randomised manner. The upper layer of the soil-sand-mixture was discarded from the analysis in order to exclude soil that was influenced by external effects like airborne spores from bacteria or fungi. Below a subsample of soil was taken and stored at 4 °C for the subsequent measurement of soil respiration. The aboveground biomass was cut, roots were carefully taken out of the soil, washed and all plant parts and the mesh bags were dried at 40°C.

Roots and shoots were weighed. Dried litter was removed from the mesh bags and the weight loss determined gravimetrically. Soil respiration was measured with the ‘MicroResp Soil Respiration System™’ using 2 g of soil per measurement and two technical replicates. CO_2_ production was measured spectrophotometrically before and after incubation of 6 hours at room temperature. After the measurements, samples were dried and soil water content was determined.

### 2.5. Statistics

We estimated the response surfaces of the measured variables along the gradient of tire particle concentration, fitting a generalized additive model (Hastie and Tibshirani, 1986; Wood, 2017). We consider fitting a generalized additive model to be a reasonable approach, because we do not know the shapes of the curves *a priori*. A spline curve was fitted based on restricted maximum likelihood (Wood, 2011), and the other parameters were set as default. We used the package “mgcv” v1.8-23 (Wood, 2018) in R v4.0.2 (R Core Team, 2020). Model performance was assessed based on adjusted R-squared, and the fitted curve shapes with the 95% Confidence Intervals (CIs). Note that, while we report p-value, we do not assess whether a result is statistically significant or not based on p-value but focus on the strength of the variable association with variability explained (R^2^), following the suggestion by the American Statistical Association (Wasserstein et al., 2019). The R script with the dataset is available for reproducibility at ‘tire-particles-in-soil’ (github repository, full link available below).

## 3. Results

The addition of tire abrasion particles negatively affected shoot and root growth. Shoot dry weight started to decrease with the lowest TP addition of 10 mg kg^−1^ and further decreased until it leveled off around 60 mg kg^−1^ (adjusted R^2^ = 0.20, p = 0.009; Fig. 1A). Root weight slightly but continuously decreased with increasing TP addition (adjusted R^2^ = 0.07, p = 0.09; Fig. 1B).

**Fig. 1:**
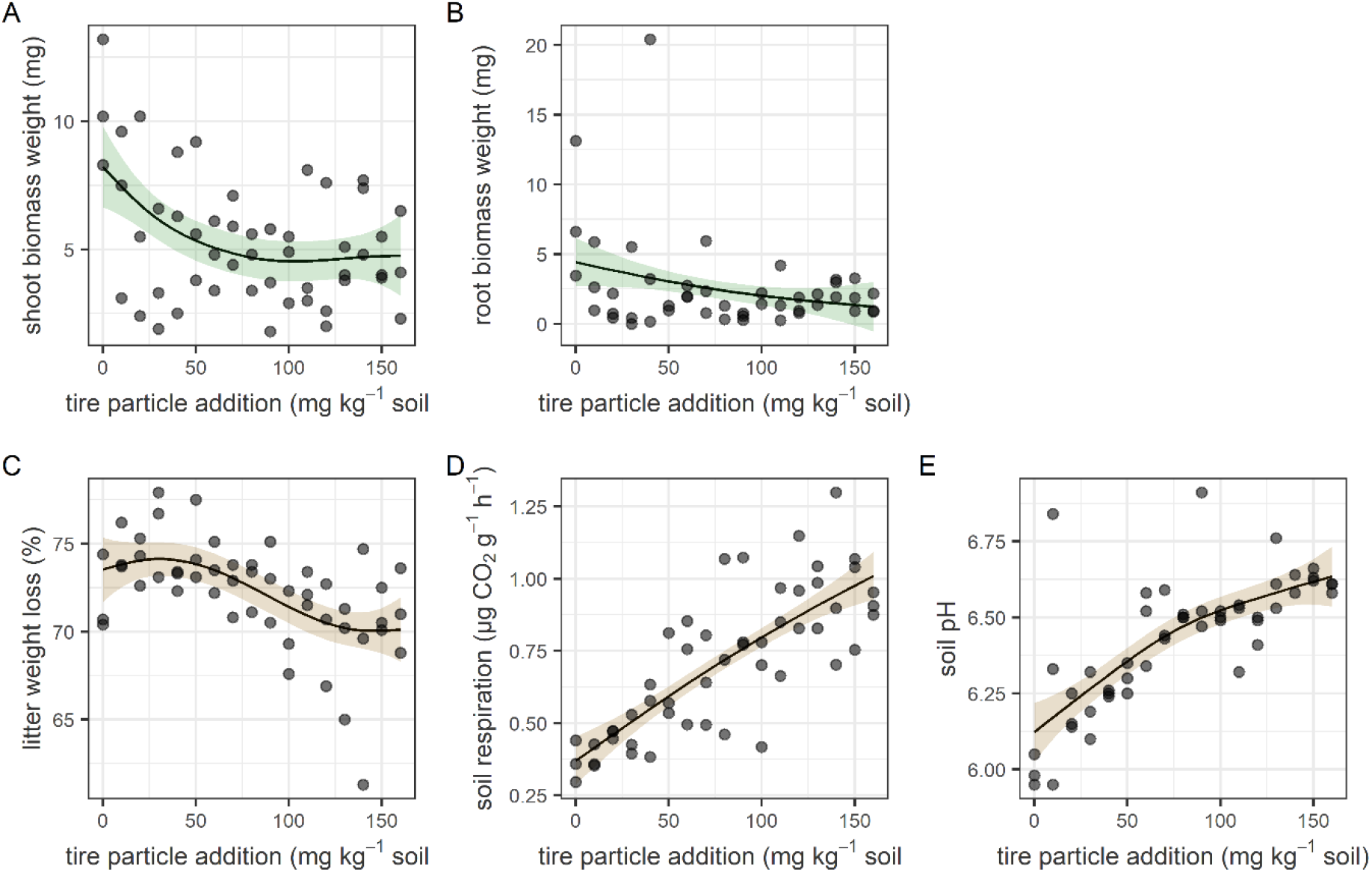
Individual data points for shoot (A) and root (B) biomass weight (mg), litter weight loss (%) (C), soil respiration (D) and soil pH (E). Lines indicate smoothed splines of response variables in response to the concentration gradient of tire particles, colored areas represent the 95% CIs.

The decomposition of the tea litter slightly increased with lower additions of TPs but at TP concentrations larger than 60 mg kg^−1^ decreased slightly with increasing concentration of TPs until an average slightly lower than that of the control was reached (adjusted R^2^ = 0.30, p = 0.0005; Fig. 1C). Soil pH increased until a TP concentration of 80 mg kg^−1^ and afterwards seemed to level off towards the end (adjusted R^2^ = 0.54, p = 6e-09, Fig. 1E). Soil respiration clearly increased with increasing concentration of added TPs (adjusted R^2^ = 0.63, p = 2e-12; Fig. 1D).

## 4. Discussion

In this study, we show that there is a direct effect of pure lab-produced TPs under controlled laboratory conditions. In previous studies Zn has been shown to be predominantly responsible for detrimental effects on plants, as this metal showed the highest increase in concentration in plant tissue, causing growth reductions, alterations of morphology and anatomy (Bowman, 1994; Schulz, 1987). According to the guidelines for safe limits for agricultural soil by the European Commission (EC, 2002) soils should have < 300 mg Zn kg^−1^, < 300 mg Pb kg^−1^, <140 mg Cu kg^−1^, <75 mg Ni kg^−1^ and <150 mg Cr kg^−1^. According to these thresholds, the tire material used for this experiment contaminated the soil with Zn with TP additions of ≥ 50 mg kg^−1^ soil. This is around the concentration level at which plant growth was not further reduced. We assume that the plants reduced their growth with starting contamination and stopped growing when Zn contamination reached a certain level in the soil. In some countries thresholds for safe limits for agricultural soils are much lower (EC, 2002), e.g. Denmark has a limit for good soil quality for Cr at 30 mg kg^−1^. Accordingly, our experimental soil was slightly contaminated with Cr and could have also impacted plant performance. Root growth slightly decreased with increasing concentration of TPs, indicating increased uptake of contaminants, especially Zn, which induced a growth reduction. Plants generally restrict metals more to root than shoot biomass as a stress-avoidance strategy to circumvent negative impacts of heavy metal pollution of the soil (Audet and Charest, 2008). This could explain the increasing effect of TPs on roots and the leveling off effect on shoots.

The availability of heavy metals depends on the pH of the soil and is higher in the lower pH range (Xian and In Shokohifard, 1989). Our experimental soil was acidic at the start of the experiment (pH 5.3), which probably facilitated their uptake. The TP addition increased the soil pH but saturated around pH 6.5 (see Fig. 1E and next section). The effect of TP on plants will differ with abiotic conditions such as soil pH, but also moisture, texture and temperature. A change in soil pH will not only affect bioavailability of heavy metals to plants but also soil processes.

The presence of TPs alters a number of biogeochemical soil parameters that can influence the decomposer community: i) TPs contain toxic components that leach from the particles, such as Zn. Zn has been shown to have harmful effects on some soil fungi and bacteria (Babich and Stotzky, 1978). ii) TPs alter bulk density and soil aeration, with consequences for water and oxygen availability. iii) TPs contain carbon, which introduces a change in resource availability iv) TPs contain CaCO_3_ as filler and thus increase the soil pH, which can influence decomposer activity and composition (Baensch-Baltruschat et al., 2020; Krishna and Mohan, 2017; Smolders and Degryse, 2002; Tao et al., 2019). v) Some components of TPs are hydrophobic (e.g. antioxidants and PAHs) (CSTEE, 2003; Wagner et al., 2018), thus potentially changing water relations in the soil. Some of these changes might have a stimulating effect on decomposers at low TP concentrations (such as improved aeration or elevated pH), while others (such as Zn contamination) have increasingly negative impacts on the decomposer community, thus cumulatively overshadowing the previous positive impacts.

At optimal soil moisture conditions, such as in our experiment with constant adjustment to 60% WHC, decreased soil bulk density and enhanced substrate aeration likely improved the conditions for microbial activity. Additionally, TPs represent a carbon source, as carbon black and the synthetic polymers contain a considerable amount of carbon (Kreider et al., 2010). Although TPs and associated particles from other sources (e.g. road abrasion) generally have a low bioavailability (Wik and Dave, 2009), some of these compounds leach from the TPs (Wagner et al., 2018) and thus are available as a resource to microorganisms and can boost their activity. However, regarding the toxicity of the particles, we assume that the toxic components only selectively affected soil fungal and bacterial species (Babich and Stotzky, 1978). Other species that are less sensitive to these pollution agents might have profited from the carbon addition. Our findings are in line with field observations where spots closer to a road showed increased soil respiration compared to spots more far from the road (Kocher et. al, 2001).

## 5. Conclusions and perspectives

In the environment TPs form heteroaggregates together with road wear particles (e.g. road abrasion particles, de-icing salts), brake ware and particles from atmospheric deposition (Charters et al., 2015). The morphology, chemical composition and potential toxicity of these ‘tire- and roadwear particles’ (TRWP) will be different from pure TPs (Baensch-Baltruschat et al., 2020), with differing impacts on the environment (Rillig et al., 2019). For example, in the presence of ions from de-icing salts, Zn forms complex ZnCl species that exert greater toxicity than Zn alone (Babich and Stotzky, 1978). However, there is a wide range of observed toxicity of tire containing particles, which can be attributed to differences in particle composition, applied concentrations, test designs and species sensitivity.

Pristine TPs are prone to degradation, resulting in an alteration of their crystalline structure, chemical composition, shape and surface texture. Degraded TRWP can release sorbed compounds and leachate over time (Councell et al., 2004; Smolders and Degryse, 2002), leading to an accumulation of harmful components in the soil. Considering the quantities of yearly produced TRWP, their persistence, and toxic potential, we assume that these particles will eventually have a significant impact on the terrestrial environment. Effects on plant diversity and soil functions (filtering, medium for plant growth, habitat for soil organisms, carbon reservoir) should be the focus of further studies.

## 6. Acknowledgements

EFL acknowledges funding from the Deutsche Forschungsgemeinschaft (LE 3859/1-1). MCR acknowledges support from an ERC Advanced Grant (694368).

## Author contributions

EFL and MCR designed the study. HLK and EFL performed the study. HLK, EF and EFL performed the lab work. MR performed the statistical analysis. EFL led the writing of the manuscript and wrote the first draft. All authors contributed to the writing of the manuscript.

## Data availability

‘Tire-particles-in-soil’. Github repository. Dataset and R code available at https://github.com/Dr-Eva-F-Leifheit/tire-particles-in-soil

## Supplementary Information

Figure S1 is available as supplementary material.

## References

Audet, P., Charest, C. (2008). Allocation plasticity and plant–metal partitioning: Meta-analytical perspectives in phytoremediation. Environmental Pollution 156:290–296. doi:https://doi.org/10.1016/j.envpol.2008.02.010

Babich, H., Stotzky, G. (1978). Toxicity of zinc to fungi, bacteria, and coliphages: influence of chloride ions. Applied and Environmental Microbiology 36:906–914

Baensch-Baltruschat, B., Kocher, B., Stock, F., Reifferscheid, G. (2020). Tyre and road wear particles (TRWP) -A review of generation, properties, emissions, human health risk, ecotoxicity, and fate in the environment. Science of the Total Environment 733:137823. doi:https://doi.org/10.1016/j.scitotenv.2020.137823

Blettler, M.C.M., Abrial, E., Khan, F.R., Sivri, N., Espinola, L.A. (2018). Freshwater plastic pollution: Recognizing research biases and identifying knowledge gaps. Water Research 143:416–424. doi:https://doi.org/10.1016/j.watres.2018.06.015

Bowman, D.C. (1994). Growth of Chrysanthemum with Ground Automobile Tires Used as a Container Soil Amendment. Hort Science 29:774–776

Brandsma, S.H., Brits, M., Groenewoud, Q.R., van Velzen, M.J.M., Leonards, P.E.G., de Boer, J. (2019). Chlorinated Paraffins in Car Tires Recycled to Rubber Granulates and Playground Tiles. Environmental Science & Technology 53:7595–7603. doi:10.1021/acs.est.9b01835

Büks, F., Loes van Schaik, N., Kaupenjohann, M. (2020). What do we know about how the terrestrial multicellular soil fauna reacts to microplastic? SOIL 6:245–267. doi:10.5194/soil-6-245-2020

Charters, F.J., Cochrane, T.A., O’Sullivan, A.D. (2015). Particle size distribution variance in untreated urban runoff and its implication on treatment selection. Water Research 85:337–345. doi:https://doi.org/10.1016/j.watres.2015.08.029

Councell, T.B., Duckenfield, K.U., Landa, E.R., Callender, E. (2004). Tire-wear particles as a source of zinc to the environment. Environmental Science & Technology 38:4206–4214. doi:10.1021/es034631f

CSTEE (2003). Opinion of the Scientific Committee on toxicity, ecotoxicity and the environment (CSTEE) on ‘Questions to the CSTEE relating to scientific evidence of risk to health and the environment from polycyclic aromatic hydrocarbons in extender oils and tyres’. European Commission Health & Consumer Protection Directorate-General, Directorate C – Public Health and Risk Assessment, Brussels

de Souza Machado, A.A. et al. (2019). Microplastics Can Change Soil Properties and Affect Plant Performance. Environmental Science & Technology 53:6044–6052. doi:10.1021/acs.est.9b01339

de Souza Machado, A.A., Lau, C.W., Till, J., Kloas, W., Lehmann, A., Becker, R., Rillig, M.C. (2018). Impacts of Microplastics on the Soil Biophysical Environment. Environmental Science & Technology 52:9656–9665. doi:10.1021/acs.est.8b02212

Ding, J. et al. (2020). Dysbiosis in the Gut Microbiota of Soil Fauna Explains the Toxicity of Tire Tread Particles. Environmental Science & Technology 54:7450–7460. doi:10.1021/acs.est.0c00917

EC (2002). Heavy Metals in Waste, Project ENV.E.3/ETU/2000/0058.

European Commission DG ENV. E3, EN 15933:2012 Sludge, treated biowaste and soil – Determination of pH.

Evangeliou, N., Grythe, H., Klimont, Z., Heyes, C., Eckhardt, S., Lopez-Aparicio, S., Stohl A., (2020). Atmospheric transport is a major pathway of microplastics to remote regions. Nature Communications 11:3381. doi:10.1038/s41467-020-17201-9

Hartmann, N.B. et al. (2019). Are We Speaking the Same Language? Recommendations for a Definition and Categorization Framework for Plastic Debris. Environmental Science & Technology 53:1039–1047. doi:10.1021/acs.est.8b05297

Hastie, T., Tibshirani, R. (1986). Generalized Additive Models. Statistical Science 1:297–310

Haward, M. (2018). Plastic pollution of the world’s seas and oceans as a contemporary challenge in ocean governance. Nature Communications 9:3. doi:10.1038/s41467-018-03104-3

Klöckner, P., Reemtsma, T., Eisentraut, P., Braun, U., Ruhl, A.S., Wagner, S. (2019). Tire and road wear particles in road environment – Quantification and assessment of particle dynamics by Zn determination after density separation. Chemosphere 222:714–721. doi:https://doi.org/10.1016/j.chemosphere.2019.01.176

Kocher, B., Siewert, C., Lorenz, M., Wolf, U. Proceedings of the 6th International Conference on the Biogeochemistry of Trace Elements; Guelph, Canada, July 2001; p 571.

Kole, P.J., Lohr, A.J., Van Belleghem, F., Ragas, A.M.J. (2017). Wear and Tear of Tyres: A Stealthy Source of Microplastics in the Environment. International journal of environmental research and public health 14. doi:10.3390/ijerph14101265

Kreider, M.L., Panko, J.M., McAtee, B.L., Sweet, L.I., Finley, B.L. (2010). Physical and chemical characterization of tire-related particles: Comparison of particles generated using different methodologies. Science of the Total Environment 408:652–659. doi:https://doi.org/10.1016/j.scitotenv.2009.10.016

Krishna, M.P., Mohan, M. (2017). Litter decomposition in forest ecosystems: a review. Energy, Ecology and Environment 2:236–249. doi:10.1007/s40974-017-0064-9

Lehmann, A., Leifheit, E.F., Feng, L., Wulf, A., Bergmann, J., Rillig, M.C. 2020. Microplastic fiber and drought effects on plants and soil are only slightly modified by arbuscular mycorrhizal fungi. Soil Ecology Letters (in press). doi.org/10.1007/s42832-020-0060-4

Leifheit, E.F., Verbruggen, E., Rillig, M.C. (2015). Arbuscular mycorrhizal fungi reduce decomposition of woody plant litter while increasing soil aggregation. Soil Biology & Biochemistry 81:323–328. doi:10.1016/j.soilbio.2014.12.003

Panko, J.M., Chu, J., Kreider, M.L., Unice, K.M. (2013). Measurement of airborne concentrations of tire and road wear particles in urban and rural areas of France, Japan, and the United States. Atmospheric Environment 72:192–199. doi:https://doi.org/10.1016/j.atmosenv.2013.01.040

Qi, Y. et al. (2018). Macro- and micro-plastics in soil-plant system: Effects of plastic mulch film residues on wheat (Triticum aestivum) growth. Science of the Total Environment 645:1048–1056. doi:https://doi.org/10.1016/j.scitotenv.2018.07.229

R Core Team (2020). R: A language and environment for statistical computing. R Foundation for Statistical Computing, Vienna, Austria. URL https://www.R-project.org/.

Rhodes, E.P., Ren, Z., Mays, D.C. (2012). Zinc Leaching from Tire Crumb Rubber. Environmental Science & Technology 46:12856–12863. doi:10.1021/es3024379

Rillig, M.C., Lehmann, A. (2020). Microplastic in terrestrial ecosystems. Science 368:1430–1431. doi:10.1126/science.abb5979

Rillig, M.C., Lehmann, A., Ryo, M., Bergmann, J. (2019). Shaping Up: Toward Considering the Shape and Form of Pollutants. Environmental Science & Technology 53:7925–7926. doi:10.1021/acs.est.9b03520

Schulz, M. (1987). Effects of ground rubber on Phaseolus vulgaris. Zeitschrift für Pflanzenernährung und Bodenkunde 150:37–41

Smolders, E., Degryse, F. (2002). Fate and Effect of Zinc from Tire Debris in Soil. Environmental Science & Technology 36:3706–3710. doi:10.1021/es025567p

Tao, J., Zuo, J., He, Z., Wang, Y., Liu, J., Liu, W., Cornelissen, J.H.C. (2019). Traits including leaf dry matter content and leaf pH dominate over forest soil pH as drivers of litter decomposition among 60 species. Functional Ecology 33:1798–1810. doi:10.1111/1365-2435.13413

Team, R.C. (2017). R: A Language and Environment for Statistical Computing. R Foundation for Statistical Computing, Vienna, Austria

van Kleunen, M., Brumer, A., Gutbrod, L., Zhang, Z. (2020). A microplastic used as infill material in artificial sport turfs reduces plant growth. Plants, People, Planet 2:157–166. doi:10.1002/ppp3.10071

Wagner, S., Hüffer, T., Klöckner, P., Wehrhahn, M., Hofmann, T., Reemtsma, T. (2018). Tire wear particles in the aquatic environment – A review on generation, analysis, occurrence, fate and effects. Water Research 139:83–100. doi:https://doi.org/10.1016/j.watres.2018.03.051

Wasserstein, R.L., Schirm, A.L., Lazar, N.A. (2019). Moving to a World Beyond “p < 0.05”. The American Statistician 73:1–19. doi:10.1080/00031305.2019.1583913

Wik, A., Dave, G. (2009). Occurrence and effects of tire wear particles in the environment – A critical review and an initial risk assessment. Environmental Pollution 157:1–11. doi:https://doi.org/10.1016/j.envpol.2008.09.028

Wood, S. (2017). Generalized Additive Models. Chapman and Hall/CRC. doi:https://doi.org/10.1201/9781315370279

Wood, S. (2018). mgcv: Mixed GAM Computation Vehicle with Automatic Smoothness Estimation. v1.8–23.

Wood, S.N. (2011). Fast stable restricted maximum likelihood and marginal likelihood estimation of semiparametric generalized linear models. Journal of the Royal Statistical Society: Series B (Statistical Methodology) 73:3–36. doi:10.1111/j.1467-9868.2010.00749.x

Xian, X., In Shokohifard, G. (1989). Effect of pH on chemical forms and plant availability of cadmium, zinc, and lead in polluted soils. Water, Air, and Soil Pollution 45:265–273. doi:10.1007/BF00283457

