## Supplemental Figure S1 for "Tire abrasion particles negatively affect plant growth even at low concentrations and alter soil biogeochemical cycling"

The following Supporting Information is available for this article:

Figure S1

**Figure S1**

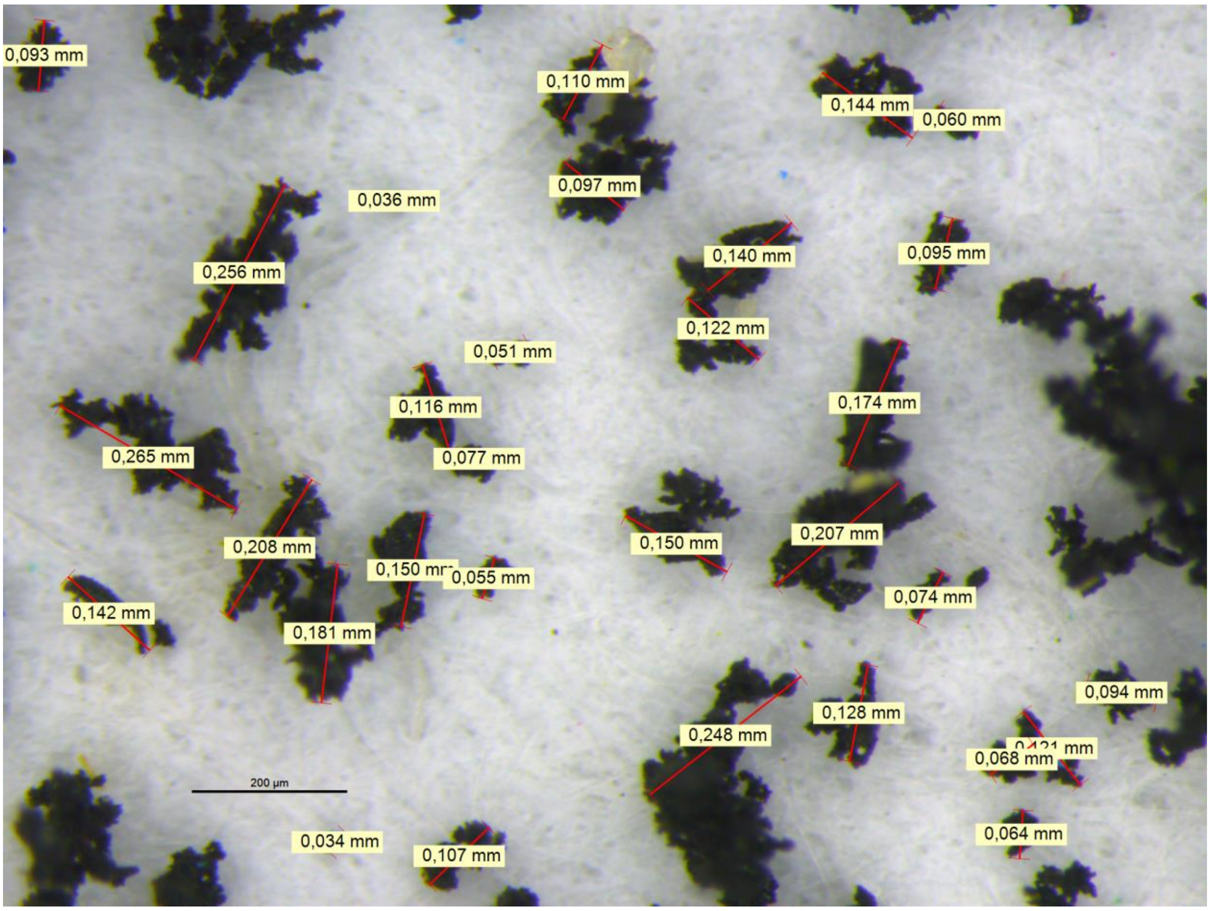


**Fig. S1**: photographic processing of tire particle size to determine average diameter and size range. Always the longest length of the particle was chosen.
